# Transforming neural activity to operate without glucose metabolism using brain-derived ketone bodies

**DOI:** 10.64898/2026.02.19.706813

**Authors:** Hafsa Yaseen, Karissa Cisneros, Rebecca Wright, Nikolaus Bueschke, Joseph M. Santin

## Abstract

The vertebrate brain is exquisitely sensitive to disruptions in glucose metabolism, and failure of adequate glucose delivery causes neurological dysfunction. Here, we discovered an animal with the capacity to defy this rule: We show that neural activity in frogs, animals with seemingly typical glucose demands, can stop metabolizing glucose by, in part, shifting to ketone bodies made exclusively within the brain after emergence from hibernation. This involves ketone body synthesis and transport from astrocytes to neurons to power synaptic transmission, along with the upregulation of gene expression that controls fatty acid catabolism and ketone body transport. Brain-derived ketone bodies also prevent decrements in activity that occur during hypoxia. These results provide insight into how frogs restart brain circuits following months of underwater hibernation when facing severe hypoxia and hypoglycemia that otherwise strongly impair neural performance in most animals. More broadly, they reveal the vertebrate brain has the capacity to serve as its own fuel reserve during the cessation of glucose metabolism, switching seamlessly to locally sourced ketone bodies while maintaining neural activity. This reframes glucose metabolism in the vertebrate brain not as a hard-wired necessity, but as a plastic trait that can in some cases be entirely abandoned.

**Significance:** The brain relies on a continuous supply of glucose from the blood to support the large energy demands of neural activity. When glucose delivery is disrupted, neural activity collapses within minutes. Here, we demonstrate that hibernation induces a large capacity for frogs, animals with seemingly normal glucose needs, to operate neural circuits without glucose metabolism, replacing it ketone bodies produced exclusively within the brain. These results reveal that a brain-derived, non-glucose fuel reserve can power neural function in the absence of glucose delivery. These findings reframe neural activity’s reliance on continuous glucose metabolism as a plastic trait, rather than a hard-wired constraint.

## Introduction

Brain activity is energetically expensive, and failure to produce ATP through aerobic respiration swiftly impairs neural function. As oxidative metabolism of glucose accounts for most of the ATP production in the brain (1), it is not surprising that defects in glucose metabolism contribute to multiple neurological issues seen in stroke, Alzheimer’s disease, Parkinson’s disease, ALS, aging, and many more (2). While fuels, like lactate and ketone bodies, can supplement some of the brain’s glucose needs under certain conditions, dietary states, or nutritional therapies, in isolation they are unable to maintain neural activity in the mammalian brain after early developmental stages, and glucose is obligatory (3–7). For example, during large shifts to brain ketone body consumption during fasting or ketogenic diets, ketone bodies fully support basal housekeeping energy needs of neurons, but account for only approximately half of activity-dependent oxidative requirements, with the remainder requiring glucose from gluconeogenesis (5). Therefore, without consistent glucose delivery, there is not enough energy to power costly physiological processes that sustain neuronal activity (6).

Although most animals obey this rule, several species, such as painted turtles, naked mole-rats, ground squirrels evolved to survive glucose and/or oxygen limitations encountered as a part of their life histories (8–10). Most amphibians are not generally counted among those strongly tolerant of brain energy stress, with neural activity having oxygen and glucose needs of an average vertebrate (11, 12). Yet, there is more than meets the eye. During aquatic hibernation, frogs have low metabolic rates and respire only through their skin while submerged in cold water. The combined effects of low metabolic rate, lack of food, and skin gas exchange cause oxygen and body glycogen stores to fall (13). This is not an immediate threat due to low energy requirements and reduced behavioral demands, but to emerge in the spring, vital neural circuits must accelerate activity in the face of severe energy limitations, where neural activity should be difficult if not impossible (12). To overcome these constraints, hibernation transforms brainstem circuits to produce activity with little energy, running for at least 3 hours during anoxia and 2 hours during oxygen and glucose deprivation (14). This large improvement during anaerobic and hypoglycemic conditions occurs by shifting to brain glycogen as likely glucose source to uphold glycolysis (14), bolstering synaptic glycolysis (15), and maintaining synaptic transmission while reducing its energetic cost (16). Therefore, a vertebrate with seemingly typical metabolic requirements can switch to local brain energy reserves to support behavior in times of stress (11).

Past studies illuminated physiological modifications that prolong neural activity during fully anaerobic conditions. However, emerging frogs are likely to be hypoxic, rather than completely without oxygen (anoxic), whereby the presence of some oxygen suggests that brain-derived fuels for aerobic ATP production may play a role. Although the brain is not generally considered to maintain large onboard fuel stores, astrocytes store two important macromolecules; glycogen that can produce lactate (17) and lipid droplets that can be converted to ketone bodies (18), two alternative fuels that when provided externally can support brain aerobic metabolism during metabolic stress (4). Therefore, we hypothesized that internal energy sources that feed aerobic metabolism may maintain neural function during aglycemic and hypoxic stress after emergence from hibernation. Here we present several lines of evidence that hibernation triggers a metabolic plasticity response that allows motor circuits to function without glucose metabolism by switching seamlessly to ketone bodies produced inside the brain.

## Results

### Neural activity drops its glucose requirement after hibernation

We assessed the function of the brainstem circuit that generate breathing, as it is a rhythmically active motor circuit critical to survival after hibernation (11). In addition, this network can be studied *ex vivo*, producing physiological motor patterns that resemble *in vivo* activity (19). *Ex vivo* tissue preparations also allow isolation of metabolic manipulations to the brain without the confounding influence from body energy stores and cerebral blood flow. To assess neural activity’s requirement for glucose metabolism, we replaced D-glucose with 2-deoxy-D-glucose (2DG-0G) under aerobic conditions with no additional fuels supplemented. With this treatment, aerobic metabolism by mitochondria remains intact, but there is no capacity for glucose metabolism because external glucose is removed and 2DG leads to the production of 2-deoxy-glucose-6-phosphate, which competitively inhibits hexokinase.

As expected (12), network activity in controls was strongly disrupted by 2DG-0G (Figure 1 A, top panel, Figure 1B). In hibernators exposed to 2DG-0G under the same conditions (Figure 1A, bottom panel), network burst rate was maintained at baseline frequency through 60 minutes and persisted significantly longer than in controls (Figure 1B). While activity slowed on average by the end of 90 minutes, all preparations produced activity that appeared outwardly normal. Exposure to 0% O2 to stop aerobic respiration quickly silenced all hibernator circuits running in 2DG-0G (Figure 1A, bottom panel). We then assessed two of the most energetically-demanding cellular processes needed to produce circuit output: excitatory synaptic transmission and action potential firing. For this, we measured AMPA-glutamatergic transmission (Figure Supplement 1) and firing activity in hypoglossal motoneurons from brainstem slices, as these neurons are part of the respiratory network (20). In controls, rhythmically evoked synaptic transmission to generate patterned input like that observed in the intact network declined steadily over 30 minutes in 2DG-0G and partially recovered upon washout (Figure 1C top panel, Figure 1D). In hibernators, synaptic transmission was maintained at baseline levels during and after 2DG-0G (Figure 1C bottom panel, Figure 1D). Firing rates and intrinsic membrane properties were maintained in 2DG-0G in both controls and hibernators (Figure Supplement 2), consistent with previous reports that these processes are energetically cheaper compared to excitatory synapses (15, 21). These results demonstrate that network activity and synaptic transmission, thought to critically rely on glucose metabolism (22), can switch to a non-glucose, aerobic energy source generated within the brain.

**Figure 1.**
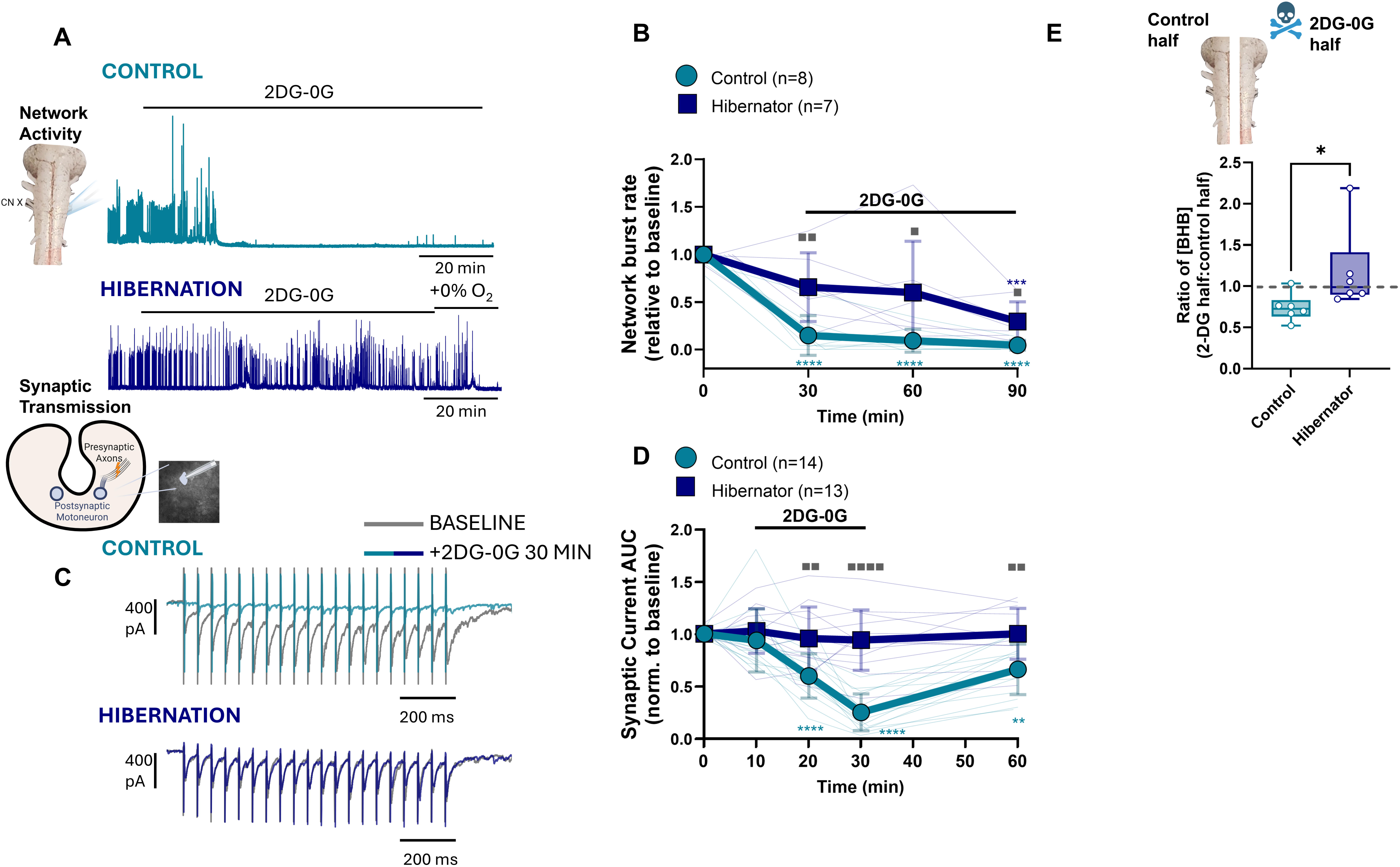
**Neural activity abandons glucose metabolism after hibernation while maintaining tissue beta hydroxybutyrate homeostasis** A) Example extracellular recordings of cranial nerve X output driven by rhythmic respiratory circuits upon replacing glucose with 5 mM 2-DG (2DG-0G) in a control (top) and hibernator in the presence of O2. After 90 minutes in 2DG-0G, O2 was removed to confirm hibernators were maintaining activity without glucose metabolism using aerobic respiration. Inset illustrates the brainstem preparation. B) Mean data of network burst rate over 90 minutes in 2DG-0G. ▪ indicates differences across groups at each time point in 2DG-0G, and * represent differences from baseline within each group. Repeated Measures Two-way ANOVA results: Time, p<0.0001, Group, p=0.0034, Time x Group, p=0.0145. **** indicates p<0.0001, ▪ p<0.05, and ▪▪ <0.01 in Holm-Sidak Multiple Comparisons tests. C) Example recordings of rhythmically-evoked AMPA-glutamate synaptic transmission onto hypoglossal motoneurons delivered for 1s at 20 Hz every 5 s following 30 minutes in 2DG-0G in controls and hibernators. Gray tracings are baseline evoked rhythmic synaptic transmission. Colors (green=control, blue=hibernator) show transmission after 30 minutes in 2DG-0G. In hibernators, it is difficult to discern the baseline because transmission was the same after 30 minutes in 2DG-0G, while in controls, activity was nearly abolished. D) Mean data of synaptic transmission (area under the curve; AUC of the transmission envelope upon rhythmic stimulation) during and after recovery from 2DG-0G. ▪ indicates differences across groups at each time point in 2DG-0G, and * represent differences from baseline within each group (control, n=14 neurons from N=6 animals, hibernators, n=13 neurons from N=6 animals). Repeated Measures Two-way ANOVA results: Time, p<0.0001, Group, p<0.0001, Time x Group, p<0.0001. ** indicates p<0.01, ****p<0.0001, ▪▪▪▪ p<0.0001, and ▪▪ p<0.01 from Holm-Sidak Multiple Comparisons tests. E) Ratio of tissue BHB concentration in the brainstem half exposed to 2DG-0G for 60 minutes relative to the control half maintained in normal conditions. Controls decreased BHB, while hibernators maintained tissue BHB. * indicates p<0.05 from Mann-Whitney U test.

### A key role for ketone bodies to support activity without glucose metabolism

While there are many potential alternative fuels, those made by astrocytes, including lactate and ketone bodies, make strong candidates. To determine the suitability of BHB (the most bioavailable ketone body) and lactate to support neural activity, we substituted either alternative fuel for glucose and assessed network output. Controls required glucose and did not function when replaced with either BHB or lactate (Figure Supplement 3A-F). Hibernators performed better than controls on each fuel but maintained activity at baseline levels with BHB while activity fell significantly with lactate. These results suggest that BHB is more effective in the absence of glucose than lactate after hibernation. To address ketone body production and consumption within the isolated brainstem, we measured tissue BHB content in two brainstem halves from each animal, one bathed in normal aCSF and the other in 2DG-0G with no supplemented alternative fuel for one hour. Absolute BHB content was not different across groups (Figure Supplement 3G). However, within the control group, tissue BHB decreased during 2DG-0G relative to the control half (Figure 1E, Figure Supplement 3G), indicating that endogenous ketone bodies are consumed when glucose is absent. Yet, as network activity and synaptic transmission fell strongly over this time (Figure 1A &1B), BHB consumption was insufficient to maintain activity. In contrast, hibernators maintained tissue BHB during 2DG-0G (Figure 1E, Figure Supplement 3G) while also sustaining neural activity and synaptic function (Fig 1A & 1B). Altogether, these results suggest that hibernation improves the ability to run on BHB without glucose and that it may be synthesized internally at a rate sufficient to prevent net decreases while being consumed to fuel ongoing activity.

### A shift to brain-derived ketone bodies power neural activity without glucose metabolism

To test the hypothesis that hibernators power neural activity with endogenously produced ketone bodies, we measured synaptic transmission after blocking brain ketone body synthesis using 2.5 μM hymeglusin, a compound that selectively inhibits HMG-CoA synthase for the production ketone bodies (23–27). In controls, synaptic transmission in 2DG-0G fell to a similar degree with or without hymeglusin by 30 minutes (Figure 2A, 2B, 2G). In hibernators, synaptic transmission was reduced in 2DG-0G in the presence of hymeglusin, with weakened evoked transmission by 30 minutes compared to hibernators without hymeglusin (Figure 2C, 2D, 2G). To verify this outcome occurred by interfering with ketone body synthesis, we performed a rescue experiment (Figure 2E-F). After 30 minutes in hymeglusin+2DG-0G in hibernators, providing 5 mM BHB over the next 30 minutes increased transmission (Figure 2E), while transmission continued to decline in hymeglusin+2DG-0G without BHB over the 60 minute period (Figure 2F, 2H). The requirement for ketone body synthesis to maintain synaptic activity was mirrored in intact rhythmic networks from hibernators (Figure Supplement 4A-D). In this way, hymeglusin led to reduced motor amplitude of respiratory motor bursts after 90 min of 2DG-0G and also a shift to abnormal, seizure-like motor bursts, suggestive of a loss of network homeostasis that occurs during energy stress (16). These responses were reversed upon exogenous BHB exposure in the presence of hymeglusin (Figure Supplement 4E-G). When glucose metabolism was intact under baseline conditions, hymeglusin did not affect synaptic transmission (Figure Supplement 5A), suggesting the loss of glucose metabolism initiates ketone body production to preserve synaptic function. Interestingly, when assessing neuronal excitability, hymeglusin exposure with glucose present increased firing rates only in hibernators with no effect on membrane potential (Figure Supplement 5B-D), pointing to additional influences of HMG-CoA synthase activity on intrinsic excitability at baseline. Overall, these results indicate that active ketone body synthesis within the brainstem sustains synaptic transmission without glucose metabolism after hibernation.

**Figure 2.**
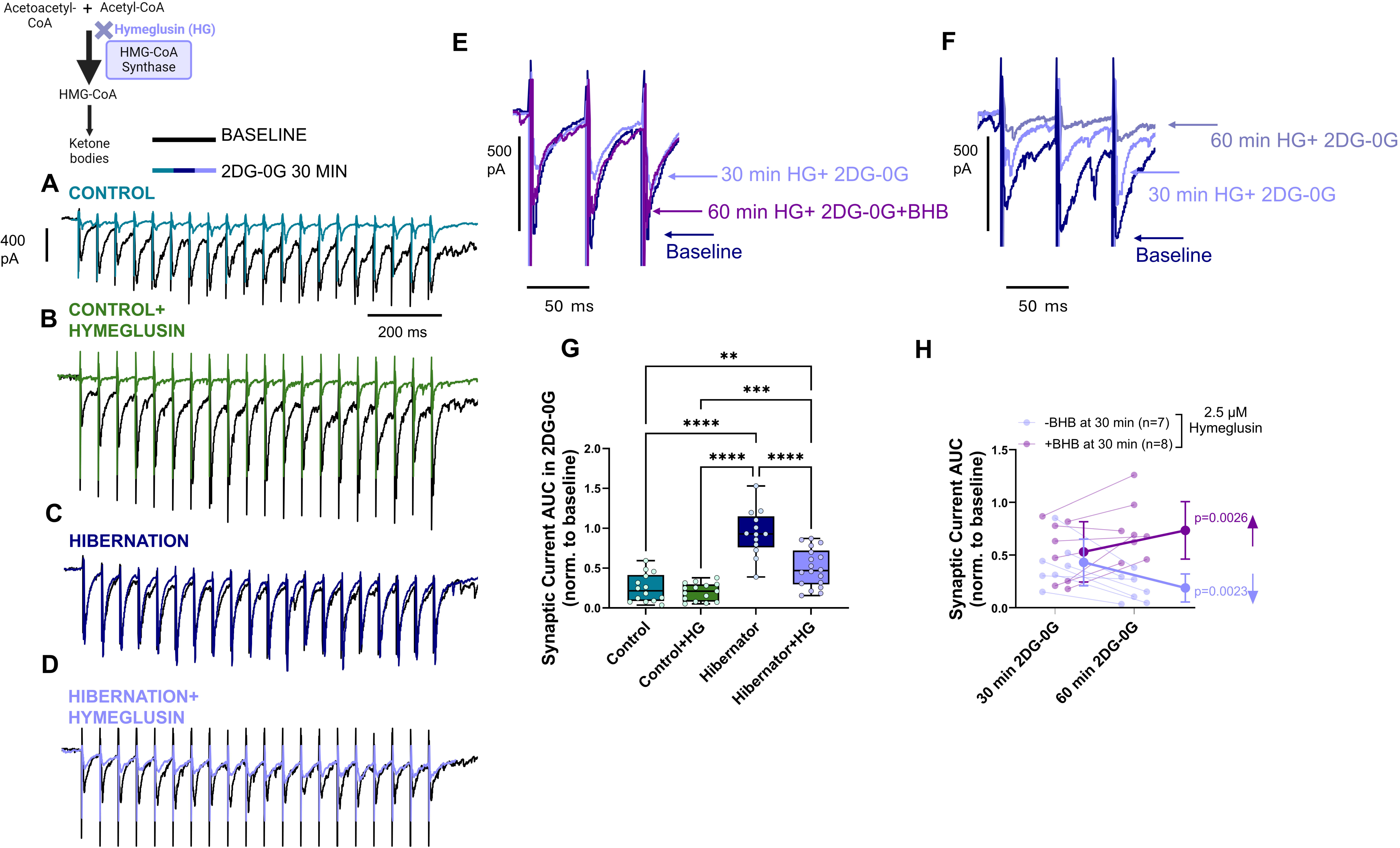
Brain ketone body synthesis maintains synaptic transmission without glucose metabolism after hibernation. A-D) Example recordings of rhythmically-evoked synaptic transmission in brainstem slices at baseline and after 30 minutes in 2DG-0G in the presence or absence of 2.5 μM hymeglusin, an inhibitor of HMG-CoA synthase, the enzyme required to produce BHB (colors). In controls, hymeglusin had no influence on the failure of synaptic transmission in 2DG-0G. After hibernation, robust synaptic transmission in 2DG-0G was reduced in the presence of hymeglusin, as indicated by smaller evoked synaptic currents during the stimulus train (light purple). E-F) Example recordings of zoomed in rhythmically-evoked synaptic transmission in hibernator brainstem slices at baseline, after 30 minutes in hymeglusin + 2DG-0G, and then either 60 minutes in hymeglusin + 2DG-0G(E) or 60 minutes in hymeglusin + 2DG-0G but with 5 mM BHB added at 30 minutes (F). By 60 minutes in 2DG-0G without BHB synthesis, transmission falls to near zero, while adding BHB at 30 minutes recovers toward baseline levels. G) Mean data of the change in synaptic transmission (area under the curve; AUC of the transmission envelope upon rhythmic stimulation) after minutes of 2DG-0G with and without hymeglusin (HG). One-way ANOVA, p<0.0001. **** indicates p<0.0001 and **p<0.01 from Holm-Sidak Multiple Comparisons Test. Control, n=14 neurons from N=6 animals. Control+HG, n=15 neurons from N=5 animals. Hibernator, n=13 neurons from N=6 animals. Hibernator+HG, n=18 neurons from 8 animals. H) Mean data showing the effect of 60 minutes in 2DG-0G in the presence of hymeglusin with BHB (dark, +BHB) or without BHB (light, -BHB) added at 30 minutes. Data points are normalized to the initial baseline of each cell. Repeated Measures two-way ANOVA results: Group x time interaction, p<0.0001, group, p=0.0165, time, p=0.6350. p values are from Holm-Sidak Multiple Comparisons Test, where transmission fell between 30 and 60 minutes without BHB while it increased when BHB was supplemented.

These results demonstrate that endogenously produced ketone bodies support synaptic activity without glucose metabolism. However, it remains unclear whether hibernation leads only to an enhanced capacity to produce ketone bodies, or whether it also bolsters the capacity of synapses to import and/or metabolize them. To address this, we added BHB during 2DG-0G in controls. As control brains have a lower biosynthetic capacity than hibernators without glucose (Fig 1E), this addresses their ability to import and use ketone bodies provided externally. In controls, providing exogenous BHB in 2DG-0G did not sustain synaptic transmission, matching controls that were not provided with supplemental BHB (Figure Supplement 6). Collectively, these results show that control frogs neither meaningfully produce nor consume BHB to support network activity. Therefore, hibernators gain both the ability to internally generate ketone bodies and use them as fuel to replace glucose metabolism.

### A role for astrocytes

Brain Ketone body synthesis is thought to involve fatty acid oxidation and subsequent production in astrocytes, as these cells have a metabolic profile comparable to liver hepatocytes (18). To test for the role of astrocytes, we incubated slices in 1 mM L-α-aminoadpic acid (AAA) for 2 hours, a treatment that disrupts the function of astrocytes in various species, including frogs (28–30). Supporting previous work that incubation with AAA induces stress and death reasonably selectively in astrocytes, acute exposure to AAA did not impair synaptic transmission (Figure Supplement 7A), and the 2 hr incubation did not affect neuronal membrane properties (Figure Supplement 7B). We compared synaptic transmission after 30 minutes in 2DG-0G in slices that were maintained in aCSF or those in aCSF that included AAA. During 2DG-0G, synaptic activity in controls was reduced to a similar extent with or without AAA (Figure 3A, 3B, 3E). In hibernators, however, 2DG-0G reduced the typical improvement in synaptic transmission after AAA treatment, with synaptic activity falling to ∼44% of baseline by 30 minutes (Figure 3C, 3D, 3E). To link astrocytes to a role in ketogenesis (Figure 2), we performed the AAA incubation in hibernators and then measured synaptic transmission while blocking ketone body synthesis with hymeglusin. If AAA exposure occluded the actions of hymeglusin, this would provide evidence for ketone body production within astrocytes. However, an additive response would suggest hymeglusin acts on cells beyond astrocytes. Combining AAA and hymeglusin produced a reduction in transmission during 2DG-0G that matched AAA alone, indicating astrocytes are the probable source of local ketone bodies. These results demonstrate that improvements in synaptic activity without glucose involves ketone body synthesis in astrocytes.

**Figure 3.**
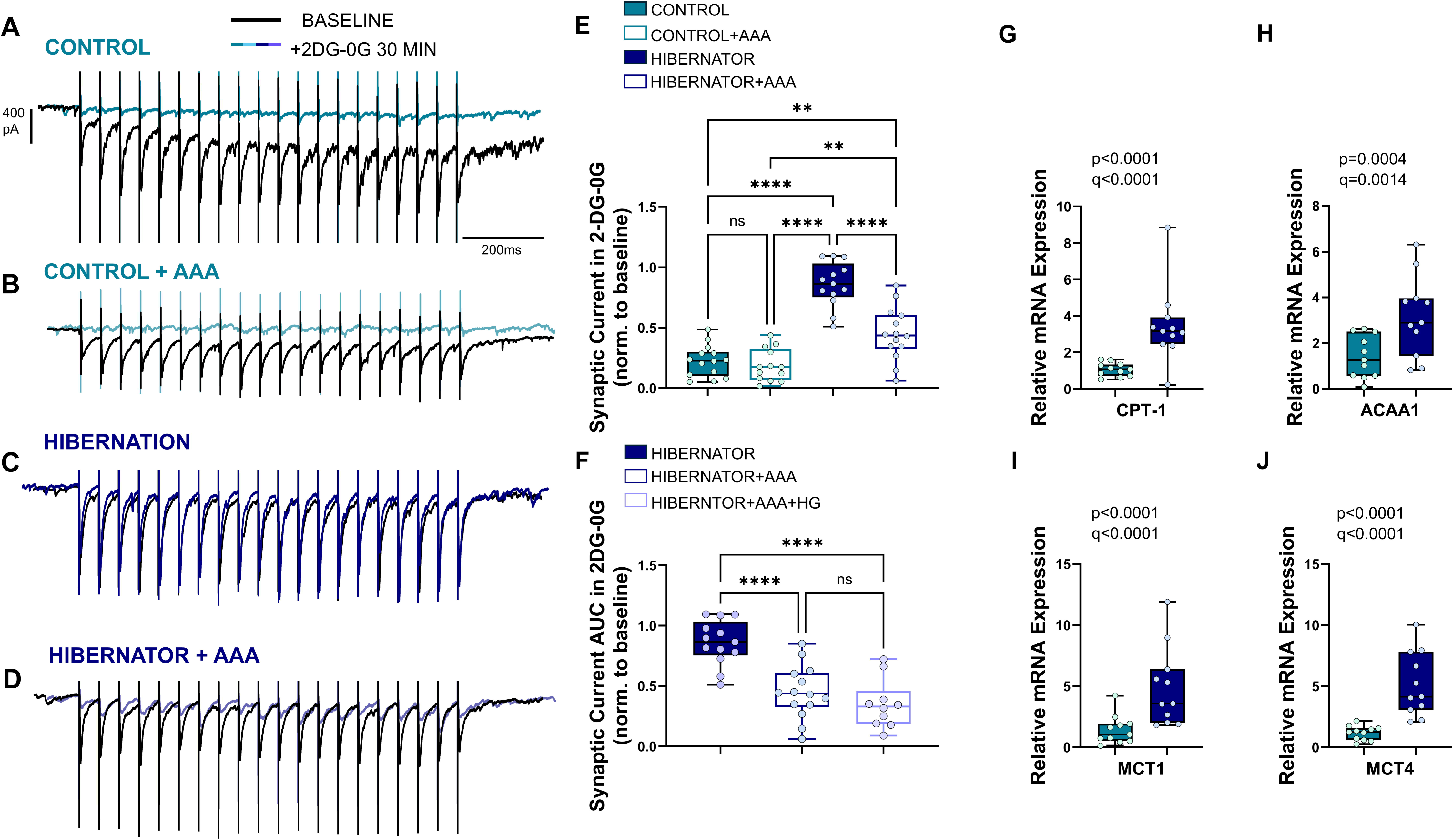
**Astrocytes contribute to synaptic activity without glucose metabolism and are associated with changes in gene expression for fatty acid metabolism and BHB transport** A-D) Example recordings of rhythmically-evoked synaptic transmission in brainstem slices in controls and hibernators with and without exposure to 1 mM L-α-aminoadpic acid (AAA) to disrupt astrocytes. Transmission in controls fell from baseline (black) in 2DG-0G with or without intact astrocytes (A and B). Hibernators maintained transmission in 2DG-0G with astrocytes intact (C), but had reduced transmission when astrocytes were disrupted by AAA (D). E) Mean data of the change in synaptic transmission (area under the curve; AUC of the transmission envelope upon rhythmic stimulation) after 30 minutes of 2DG-0G with and without astrocyte disruption by AAA. One-way ANOVA, p<0.0001. **** indicates p<0.0001 and **p<0.01 from Holm-Sidak Multiple Comparisons Test. Control, n=14 neurons from N=6 animals. Control+AAA, n=13 neurons from N=7 animals. Hibernator, n=13 neurons from N=5 animals. Hibernator+AAA, n=14 neurons from 6 animals. F) Mean data of the change in synaptic transmission (area under the curve; AUC of the transmission envelope upon rhythmic stimulation) after 30 minutes of 2DG-0G with and without astrocyte disruption by AAA and AAA+ 5 µM hymeglusin. One-way ANOVA, p<0.0001. **** indicates p<0.0001 from Holm-Sidak Multiple Comparisons Test. Hibernator, n=13 neurons from N=6 animals. Hibernator+AAA, n=14 neurons from 6 animals. Hibernator+AAA+HG, n=10 neurons from 5 animals. G-J) Increased mRNA expression for candidate genes involved in ketone body production and transport in brainstem tissue. CPT-1 (carnitine palmitoyl transferase 1), ACAA1 (acetyl-CoA acyltransferase 1), MCT1 (monocarboxylase transporter 1), MCT4 (monocarboxylase transporter 4). p values in plots are from two-tailed unpaired t test. q values are from the two-stage linear step up procedure of Benjamini, Krieger,and Yekutieli to control for the false discovery rate. N=11 per group for each gene.

An enhanced capacity of the brain to generate and use ketone bodies for fuel could relate to alternations in fatty acid breakdown, ketone body synthesis, ketone body catabolism, and/or transport. Therefore, we assessed mRNA expression of several candidate genes involved in these processes using quantitative real-time PCR in brainstem tissue homogenates. Brainstems from hibernators increased the mRNA expression of genes involved fatty acid metabolism that are thought to exist largely in astrocytes: Carnitine palmitoyl transferase I (CPT-1)(31) and peroxisomal thiolase that perform the last step of beta oxidation (ACAA2: enzyme that converts acyl-CoA into acetoacetyl-CoA)(32) (Figure 3 G-H). In addition, monocarboxylate transporters (MCT) 1 and 4, which transfer BHB across the cell membrane, were upregulated, where MCT4 is a low affinity transporter primarily expressed in astrocytes (33) (Figure 3 I-J).

Candidate genes involved in ketone body synthesis and catabolism were unaltered, along with several others for fatty acid metabolism and the neuron-specific MCT2 (Figure Supplement 8). While these results do not exclude the possibility of post-transcriptional regulation on these cellular processes, and do not explicitly isolate gene expression to specific cell types or proteins to sub-cellular compartments, they point to a brainstem-wide transcriptional program favoring fatty acid breakdown and BHB transport out of astrocytes, consistent with their role in improving synaptic function without glucose after hibernation.

### Brain-derived ketone bodies promote activity in hypoxia

Finally, brainstem activity becomes hypoxia-tolerant when frogs emerge from hibernation, where they maintain activity during tissue hypoxia and anoxia (14, 15). This results from enhanced glycolytic support and reduced aerobic requirements for neural activity (14, 15). However, BHB may modestly improve ATP yield per unit of oxygen consumed compared to glucose, particularly at low oxygen tensions, potentially by driving the mitochondrial NADH/NAD ratio higher in hypoxia (34). Therefore, the production of BHB during hypoxia could enhance energy status beyond using only glucose. To address this possibility, we measured network activity while reducing tissue pO2 with glucose metabolism intact. Corroborating previous reports (35), controls reduced network activity when tissue pO2 dropped below 30 mmHg, whereas hibernators maintained activity near baseline levels down to anoxia (Figure 4A-C). To determine if ketone body synthesis contributed to the shallow relationship between activity and pO2 in hibernators, we used hymeglusin to block ketone body synthesis and BHB replacement to rescue. In the presence of hymeglusin, activity fell from baseline at pO2 values ≤30 mmHg (Figure 4D, 4E, 4G). When we applied BHB in the presence of hymeglusin, the hypoxia-tolerant phenotype of the hibernators was restored, where activity did not significantly change from baseline down to anoxia (Figure 4F, 4G).

**Figure 4.**
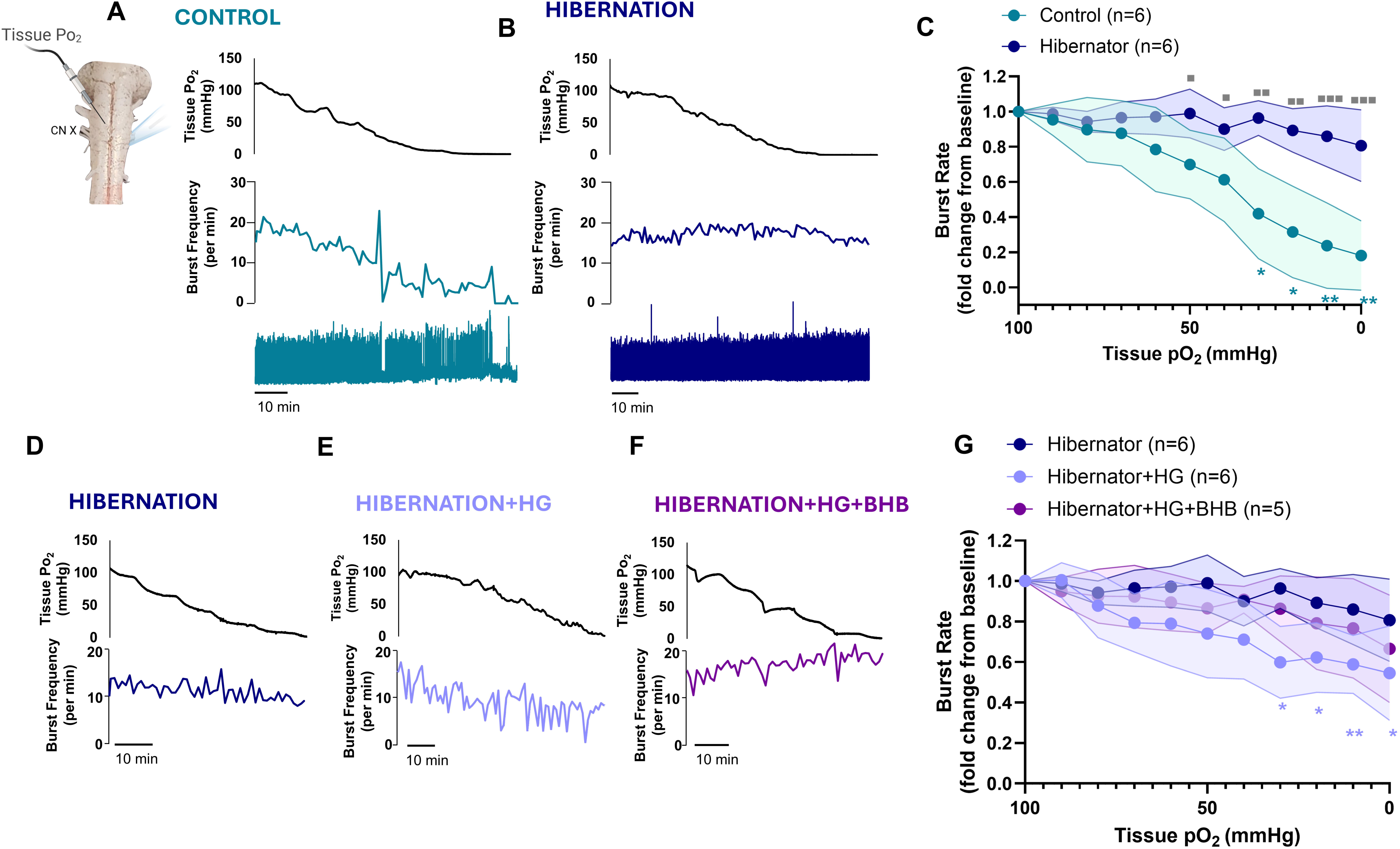
**Ketone body production contributes to improvements in network activity during hypoxia when glucose metabolism is intact** A-B) Example recordings of tissue pO2 (top), network burst rate (middle), and raw nerve activity (bottom) during graded tissue hypoxia in a control (A) and hibernator (B). In controls, as tissue pO2 progressively decreases (top panel) network burst rate also falls (middle panel). Hibernators maintain network activity as O2 falls. Inset illustrates the experimental set up used to measure network activity and tissue pO2 350 μM below the ventral surface within a component of the respiratory network. C) Mean data showing the oxygen sensitivity (change in burst rate vs. tissue pO2) in controls and hibernators. ▪ indicates differences across groups at each pO2, and * represent differences from baseline within each group. Repeated Measures Two-way ANOVA results: pO2, p<0.0001, Group, p=0.0006, pO2 x Group, p=0.0005. ** indicates p<0.01, *p<0.05, ▪▪▪ p<0.001, ▪▪p<0.01, and ▪ p<0.05 from Holm-Sidak Multiple Comparisons tests. D-F) Example recordings of tissue pO2 (top) and network burst rate (bottom) during graded tissue hypoxia in hibernators, hibernators in the presence of hymeglusin (HG), and hibernators in the presence of HG and 5 mM BHB. E) Mean data showing the oxygen sensitivity (change in burst rate vs. tissue pO2) in hibernators with no drug, 2.5 μM hymeglusin, and 2.5 μM hymeglusin + 5mM BHB. * represent differences from baseline within the hymeglusin group. Repeated Measures Two-way ANOVA results: pO2, p<0.0001, Group, p=0.0538, pO2 x Group, p=0.0182. ** indicates p<0.01, and *p<0.05, from Holm-Sidak Multiple Comparisons tests.

Finally, hymeglusin did not affect the activity vs. pO2 relationship in controls (Figure Supplement 9A), nor did it affect the apparent O2 consumption (acute change in tissue pO2 relating to O2 consumption) (35) in controls or hibernators at baseline (Figure Supplement 9B). Therefore, brain-derived ketone bodies not only power neural activity when glucose metabolism is absent, but also contribute to the maintenance of activity during hypoxia with intact glycolysis after hibernation.

## Discussion

The maintenance of neural circuit activity is widely believed to require the constant delivery and metabolism of glucose (6). Even acute impairments rapidly disrupt the physiological processes that generate neural activity (36). In contrast, we demonstrate that neural activity’s requirement for glucose is not fixed, but instead, can be remodeled and replaced with ketone bodies produced within the brain itself. Beyond early developmental periods, synapses have sizable glucose demands that cannot usually be replaced by alternative fuels, let alone those produced from brain-derived fuel stores (3, 4, 6, 7). The capacity to abandon glucose and replace it with ketone bodies generated from within the brain indicates greater metabolic flexibility than is currently appreciated for active neural circuits.

The experiments here support that frogs typically have high glucose demands and cannot maintain neural function solely using internally-derived or externally-provided ketone bodies or lactate, much like adult mammals (3, 4). However, hibernation reprograms metabolic needs such that neural activity can abandon glucose and operate on ketone bodies produced within the brain, with likely importance for restarting neural circuits as animals emerge from hibernation when oxygen and body glycogen stores are limited (11, 37). Key components of this transformation seem to involve a gained ability to: *i*. sense the loss of glucose metabolism, *ii.* increase ketone body production from products of beta-oxidation within astrocytes, *iii*. improve the capacity for release, uptake and/or metabolism of ketone bodies, and *iv.* fuel synaptic transmission while achieving network homeostasis. Mechanistically, our data support this model in the following ways: For point *i.*, blocking ketone body synthesis had no effect on baseline synaptic transmission (Figure Supplement 6A), suggesting that the loss glucose metabolism is sensed to increase ketone body synthesis to support synapses. For point *ii*., we suggest that glucose deprivation may lead to astrocytic AMP-dependent protein kinase activation, leading to the inhibition of Acetyl-CoA Carboxylase to facilitate fatty acid import into the mitochondria through CPT-1 for beta oxidation and ultimately ketone body synthesis (18), which we show to be strongly upregulated in hibernators (Figure 3G). The importance of points *iii* and *iv* is underscored by the failure of control brainstems to utilize externally provided BHB, or any that is exists internally, to support activity without glucose metabolism (Figure Supplement 3A-C, Figure Supplement 3G, Figure Supplement 6), demonstrating that the ability to import and metabolize ketone bodies is not constitutively active in frogs but must be initiated through hibernation-induced remodeling. Based on the data presented, this may occur through enhanced ketone body transport (Figure 3I-J) but may also involve post-transcriptional modifications in mitochondrial function that favor the use of ketone bodies to power synapses.

Our results support the role of brain-derived ketone bodies as a large contributor to running without glucose metabolism after hibernation in frogs. We used two manipulations that disrupt their production, hymeglusin, which inhibits HMG-CoA synthase involved in the synthesis of ketone bodies (acetoacetate and BHB), and AAA to lesion astrocytes, the principal source of ketone bodies in the brain. While our results support the role of brain-derived ketone bodies, a few possibilities may underlie the sub-cellular organization of their synthesis. In mammals, ketone body synthesis occurs primarily in mitochondria and uses HMG-CoA Synthase 2 (38). Lower vertebrates contain only the gene that codes for HMG-CoA synthase 1, which is canonically cytoplasmic rather than mitochondrial. However, HMG-CoA synthase 1 can exist in the cytosol or peroxisomes with a potential role in ketone body synthesis, as these compartments contain the full enzyme complement required to produce either BHB or acetoacetate (39–41). Furthermore, HMG-CoA synthase 1 in fish and human cancer cells can paradoxically show mitochondrial compartmentalization, activity, and a contribution to respiration (42, 43). Any of these routes, likely within astrocytes, could contribute to ketone bodies within the hibernator brain to operate during glucose impairments.

Our results also do not exclude the possibility of other internal energy sources, as manipulations that impaired ketone body production still resulted in greater synaptic transmission than controls (Figures 2&3). Recent evidence suggests that neurons may burn triglycerides during glucose deprivation (44) and energetically expensive activities like learning (45). While 2DG-0G substantially reduces glycolytic capacity, metabolites entering glycolysis downstream of hexokinase could in principle contribute to the remaining energy supply, including fructose synthesized by microglia in neurological disorders with reduced glucose metabolism (46), which improves the recovery of synaptic transmission in naked mole-rats following glucose deprivation (9). Astrocytic glycogen can be broken down to free glucose, which would fail to support neuronal metabolism under 2DG-0G conditions as it cannot bypass hexokinase inhibition.

However, astrocytic glycogen can also be mobilized as lactate, which enters glycolysis downstream of hexokinase and could in principle contribute to the remaining energy supply under 2DG-0G conditions (47). Notably, we show that BHB is a more effective fuel for maintaining neural activity than lactate after hibernation (Figure Supplement 3D-F), suggesting that even if astrocytic glycogen-derived lactate contributed under 2DG-0G conditions, it cannot fully account for the hibernator advantage. One recent study demonstrated that dopaminergic neurons contain glycogen at their terminals, providing glucose-6-phosphate derived from glycogenolysis downstream of hexokinase to sustain synaptic function (48); whether a similar mechanism operates in brainstem motor circuits studied here remains unknown. Overall, while the contributions of ketone bodies and other endogenous energy sources require further exploration, our results demonstrate a large capacity for ketone bodies synthesized within the brain to power neural activity without glucose metabolism after hibernation.

Brain-derived ketone bodies also contributed to the improvement in neural activity during hypoxia. Improved motor function in hypoxia/anoxia after hibernation was previously interpreted to result from enhanced glycolytic capacity and a lower energetic cost of activity (15, 35). It should be noted that hypoxia experiments were performed with glucose metabolism intact, and therefore, it does not exclude a contribution from glycolytically produced pyruvate to sustaining function under hypoxic conditions.

However, the requirement for ketone body synthesis to maintain activity during hypoxia, and its rescue by exogenous BHB (Figure 4D-G), suggests that brain-derived BHB confers a specific energetic advantage at low tissue pO2 beyond what glucose alone can provide. In addition to functioning as an energy source, BHB also has diverse roles in cell signaling which may contribute to this response, where it can bind to surface receptors and acutely influence neurotransmitter and neuromodulatory systems, which in principle, could also promote excitability (49). Indeed, hymeglusin, and presumably a reduction in endogenously produced ketone bodies, increased neuronal excitability when glucose metabolism was intact only in hibernators (Figure Supplement 5D). This is consistent with the ability for BHB to stimulate G-protein coupled receptors to alter neuronal excitability independent from its roles in energy metabolism (49). Regardless of the specific energetic or signaling pathways, we demonstrate that circuits can exploit locally-sourced ketone bodies to promote activity during hypoxia in addition to serving as an important energy source when glucose metabolism is absent.

These findings have general implications for understanding energetic plasticity in neural circuits. While the adult brain is thought to be constrained by glucose metabolism, it appears that some circuits are equipped with latent states that afford greater flexibility to run on internal, non-glucose fuels than is presently appreciated. Part of this resilience involves events that reprogram neural circuits to generate, transport, and metabolize ketone bodies as a fuel for synapses during glucose deprivation. A previously unknown capacity to stop glucose metabolism and replace it with locally sourced ketone bodies, and likely other endogenous fuels, is especially timely, as many neurological disorders are caused by or associated with impairments in glucose metabolism, often with ketone body metabolism left largely intact (1, 2). While there are idiosyncrasies across various animal models and humans, at a fundamental level, many of the energetic rules are conserved (50). Thus, these results provide a path to understanding how to transition active neural circuits away from glucose and replace them with locally-derived alternatives without severely compromising function. The metabolic transformation described here likely represents an acclimation response to the overwintering environment, suggesting a time component to its development.

Understanding the regulatory processes that control these state transitions has the potential to inform new strategies that protect neural activity in a wide-range of disease states associated with compromised glycolysis.

## Materials and Methods

### Animals

All experiments performed were approved by the Animal Care and Use Committee (ACUC) at the University of Missouri (protocols #39264 and #65623). Adult American Bullfrogs, *Aquarana catesbeiana*, were assigned to one of two groups: Controls and or cold-acclimated hibernators. Hibernated frogs were kept in plastic tanks in temperature control incubators for > 1 week before temperature was lowered in stepwise manner to 4°C over 7 days in a walk-in temperature-controlled environmental chamber. Once water temperature reached 4°C, air access was blocked using a plastic screen placed in the tank. After at least 3 weeks of submergence, experiments commenced.

### Neurophysiology recordings

Rhythmically active brainstem-spinal cords or brainstem slices containing hypoglossal motoneurons were generated as previously described (14,15). For experiments addressing the requirement of glucose metabolism for activity of either preparation glucose was removed from the aCSF, and 5 mM 2-deoxy-D-glucose was added (2DG-0G). Alternative fuel substitution or inhibitors were added as described in the text.

### Tissue BHB measurement and real-time quantitative PCR

Tissue BHB was measured in brainstem halves from each animal group, either exposed to control aCSF (baseline) or 2DG-0G for 60 minutes. Following this experiment both halves from each animal was assessed for BHB content, following the manufacturer’s instructions for the BHB colorimetric assay kit from Cayman Chemical following the manufactures instructions (kit #700190, Ann Arbor, MI, USA). The assessment of candidate genes was performed using standard approaches used by our lab group (16).

Complete and detailed methods for all experiments and protocols are presented in the Supplementary Material file.

## Acknowledgements

This work was supported by the following grants: National Science Foundation-Award # 2515635 (to JMS) and National Institutes of Health-R01NS114514 and R01NS144063 (to JMS). The authors thank Jose Viteri for providing comments on a draft of this manuscript.

