## Supplemental methods and figures 1-9 for "Transforming neural activity to operate without glucose metabolism using brain-derived ketone bodies"

#### Supplemental Materials for

### Brain-derived ketone bodies can replace glucose to power neural function

**Detailed Material and Methods**

#### Animals

All experiments performed were approved by the Animal Care and Use Committee (ACUC) at the University of Missouri (protocols #39264 and #65623). Adult American Bullfrogs, *Aquarana catesbeiana*, of undetermined sex were purchased from Rana Ranch (Twin Falls, Idaho, USA) and Niles Biological (Sacramento, CA, USA). Controls and cold-acclimated hibernators were maintained as previously described (1, 2). We refer to the cold-acclimated group here as "hibernators," as cold-submergence in ranid frogs leads to metabolic suppression beyond that which can be explained by reduced body temperature alone, characteristic of hibernation in vertebrates(3). Briefly, animals were kept in plastic tubs with dechlorinated tap water at room temperature bubbled with room air. Control frogs were acclimated to these conditions for at least a week after arrival at the laboratory before experiments and had access to wet and dry areas in the tanks. Pellet food was provided once per week and was eaten ad libitum. Hibernated frogs were kept in plastic tanks in temperature control incubators for > 1 week before temperature was lowered in stepwise manner to 4°C over 7 days in a walk-in temperature-controlled environmental chamber. Once water temperature reached 4°C, air access was blocked using a plastic screen placed in the tank. After at least 3 weeks of submergence, experiments commenced.

#### Tissue Preparations for Physiology Experiments

##### *Brainstem-spinal cord preparation and data processing*

Rhythmically active brainstem-spinal cords were generated as previously described(1, 2) In summary, frogs were anesthetized with isoflurane (1 ml per liter, v/v) until loss of the toe pinch reflex and then rapidly decapitated with a guillotine. The brainstem and rostral portion of the spinal cord were then dissected from the head while submerged in cold artificial cerebrospinal fluid (aCSF; concentrations in [mM]: 104 NaCl, 4 KCl, 1.4 $MgCl_2$, 7.5 glucose, 40 $NaHCO_3$, 2.5 $CaCl_2$ and 1 $NaH_2PO_4$, bubbled with 98.5% $O_2$, 1.5% $CO_2$; pH ≈ 7.85). Once exposed, the forebrain was pithed. The brainstem-spinal cord was removed with the cranial nerve roots intact and placed ventral side up in a 6 ml chamber superfused with gased aCSF. All experiments in controls and hibernators were performed at room temperature (20-22 °C). Flow rate of the superfusion was maintained at 6 ml per minute with a peristaltic pump (Ranin Rabbit 4 channel head, Mettler-Toledo Rainin, Oakland, CA, USA).

To monitor output from the respiratory motor network, we recorded activity of cranial nerve X with custom made glass suction electrodes. Nerve activity was amplified 1000X (A–M Systems Model 1700, A–M Systems, Carlsborg, WA, USA), filtered (10 Hz – 5 kHz), and digitized with a Powerlab 8/35 (ADInstruments, Colorado Springs, CO, USA). For quantification, amplified nerve output was integrated and rectified online with a T of 100 ms using the LabChart software (ADInstruments, Colorado Springs, CO, USA). Preparations were then given 2-3 hours to stabilize before experiments. For experiments addressing the requirement of glucose metabolism for activity of the preparation (controls, N=8, hibernators, N=7), glucose was removed from the aCSF, and 5 mM 2-deoxy-D-glucose was added (2DG-0G) and left for 90 minutes. To demonstrate the requirement for aerobic respiration in the maintenance of activity in 2DG-0G, after 90 minutes the aCSF was bubbled vigorously with 98.5% $N_2$/1.5% $CO_2$, a treatment that causes tissue anoxia under the conditions of our experiments (4). For experiments that assessed the ability of the network to run on beta hydroxybutyrate (BHB) or lactate with no glucose (control, N=8; hibernator, N=7), glucose was removed and substituted with equimolar BHB and 15 mM lactate. We doubled the lactate concentration compared to glucose because one glucose molecule would give rise to two lactate molecules, and previous work indicates that equimolar lactate cannot maintain activity (5). However, as pointed out by Hollnagel*, et al.* (5), we acknowledge that glucose, BHB, and lactate used here are unlikely to be isocaloric, as each produces differing amounts of ATP, and has different diffusion rates within the tissue, transport kinetics across the cell membrane, and distinct processes for intracellular metabolism.

In experiments that address the influence of hymeglusin on longevity of the hibernator neural activity during 2DG-0G, the protocol was as follows. After the baseline period in aCSF, 2.5 μM hymeglusin was added to the aCSF and superfused the preparation for 40 minutes (n=6). In naïve hibernators (n=6), no drug was added. Then 2DG-0G was applied for 90 minutes. At the end of the 90 minute period of 2DG-0G+hymeglusin, 5 mM BHB was added to the aCSF. Finally, we performed a set of experiments where oxygen in the bathing solution was gradually decreased while measuring the tissue partial pressure of $O_2$ ($pO_2$). For this, we measured tissue $pO_2$ as described recently by our group (4). To standardize the initial $pO_2$ across experiments, the electrochemical sensor was inserted into brainstem until 100 mmHg was reached. This protocol was first done in controls (n=6) and hibernators (n=6). In addition, 2 addition groups of hibernators were included: 2.5 μM hymeglusin (n=6) and 2.5 μM hymeglusin+ 5 mM BHB (n=5). Finally, hymeglusin was applied to control preparations, as well (n=6). Hymeglusin is also likely to inhibit HMG-CoA synthase involved in cholesterol synthesis; however, we are not aware of mechanisms involving the acute loss of cholesterol synthesis that would explain rapid disruption of neuronal homeostasis only in hibernators upon energy stress and rescue by ketone bodies.

For processing of nerve activity data, burst rate of activity related to breathing was counted over ten minute sampling periods and presented as bursts per minute. Breathing-related neural activity has a characteristic shape of “smooth” rising and falling phases that occur over ~1s and is the default activity mode of this preparation (Supplemental Figure 4, inset). In cases where burst amplitude of breathing-related activity was measured, the Peak Analysis module of LabChart Analysis software was used to measure the height, from the baseline to the burst peak (ADInstruments, Colorado Springs, CO, USA). Non-respiratory bursts, which are indicative of a loss of homeostasis during energy stress(2), were identified by their long and/or large amplitude relative to breathing-related bursts and longer duration (ranging from 5 to 20 seconds) (Supplemental Figure 4, inset) and presented as bursts in 10 minutes. Finally, sensitivity of the respiratory burst activity to decreases in tissue $pO_2$ were presented as % change from baseline as a function of tissue $pO_2$. Changes in $pO_2$ during fixed bath oxygen relate to changes in tissue oxygen consumption (4). To assess the potential for changes in oxygen consumption by hymeglusin application, we measured tissue oxygen after hymeglusin was applied.

*Hypoglossal motoneuron-containing slice preparation, patch clamp electrophysiology, and data processing*

Tissue slices containing hypoglossal motoneurons for patch clamp electrophysiology experiments were generated following previously established protocols (6, 7). Briefly, the brainstem-spinal cord was dissected as described for the brainstem-spinal cord preparation but instead of recording nerve activity, was sectioned using a Campden Vibrating Microtome (7000smz, Campden Instruments; Lafayette, IN, USA). For this, the brainstem-spinal cord was superglued to an agar block and then cut into 300 µM-thick sections. Slices containing the hypoglossal motor pool were given at least 1 hr to recover from the slicing procedure, transferred to a 0.5 ml chamber, and fixed to the chamber with a nylon grid for patch clamp electrophysiology experiments. The chamber was situation in a fixed stage microscope for imaging of the slice (FN1, Nikon Instruments Inc., Melville, NY, USA). Oxygenated aCSF was fed through the chamber with a gravity-fed superfusion system at ~1.5 ml per minute.

Hypoglossal motoneurons in the rostral part of the motor pool were identified anatomically based on their dorso-medial position relative to the 4$^{th}$ ventricle and large cell body size (>20 µM diameter along the shortest axis) as previously described(7). Slices were imaged using a Hamamatsu ORCA Flash 4.0LT sCMOS camera (Hamamatsu Photonics, Hamamatsu City, Japan) interfaced with NIS elements software (Nikon). Hypoglossal motoneurons in the rostral region were targeted as they are part of the buccal force pump for ventilation in anuran amphibians(8, 9) and therefore play a role in breathing and other orofacial behaviors that are critical to survival(10, 11). Once identified, motoneurons were approached by applying positive pressure through the glass microelectrode for patch clamp recording. Electrophysiological signals were amplified by an Axon Instruments 200B patch clamp amplifier and digitized with a Digidata 1550 digitizer (Molecular Devices). Microelectrodes were fabricated with a Sutter Instrument P-1000 micropipette puller (Novato, CA, USA) and had resistances of 5-6 MΩ when filled with the pipette filling solution (in mmol): 110 potassium gluconate, 2 $MgCl_2$, 10 Hepes, 1 $Na_2$-ATP, 0.1 $Na_2$-GTP and 2.5 EGTA, pH ~ 7.2 with KOH. As the motoneuron was approached, debris was cleared by the positive pressure and when an indentation was observed in the center of the cell body, positive pressure was removed, increasing the resistance between the microelectrode and the cell membrane. Negative pressure was then applied until a seal ≥1G was observed and was then broken using rapid negative pressure by mouth to obtain whole-cell access.

After obtaining whole-cell access, neurons were maintained in voltage-clamp mode at a holding voltage of -80 mV. Excitatory postsynaptic currents (EPSCs) carried by AMPA-glutamate receptors via stimulation of solitary tract (ST) axons(12) were maximally evoked using an AM-Systems isolated pulse stimulator (Model 2100, , A–M Systems, Carlsborg, WA, USA) with bipolar tungsten steroelectrodes (WE3ST0.1B10, Microprobes for Life Sciences, Gaithersburg, MD, USA) AMPA-glutamatergic identity of these synapses was determined by applying 20 μM DNQX, which eliminated evoked tranmission. Maximal synchronous EPSC amplitudes were determined by increasing the stimulus current until asynchronous vesicle release was observed (EPSCs that trailed after the synchronous EPSC and were larger than spontaneous EPSCs). The stimulus intensity was then reduced to maintain the largest possible synchronous EPSC amplitude without asynchronous activity. For experiments in this study, the main protocol was as follows. The amplifier was switched to current clamp mode to measure membrane properties (membrane potential, firing rate during stepwise current injection, and input resistance). The amplifier was switched back to voltage clamp with a holding potential of -80 mV and axons were stimulated rhythmically at maximum intensity (20 Hz, 1s duration, every 5 s) to simulate the ongoing rhythmic input to these neurons. After 5 minutes of baseline stimulation, 2DG-0G was applied for 30 minutes during rhythmic stimulation. Every 10 minutes, rhythmic stimulation was paused, and the amplifier was switched back to current clamp mode to assess membrane properties. After assessing membrane properties, the amplifier was switched back to voltage clamp mode and rhythmic synaptic stimulation resumed for the next 10 minute interval.

In some experiments (control, n=14; hibernator, n=13), glucose metabolism was restored for the next 30 minutes during rhythmic synaptic stimulation. In a different set of experiments, controls (n=15) and hibernator (n=18), this protocol was performed in the presence of 2.5 μM hymeglusin. For this, after entering whole-cell mode, baseline membrane properties were determined in current clamp, and then measured rhythmic trains of EPSCs in voltage clamp for 5 minutes. Hymeglusin was applied for 10 minutes during rhythmic synaptic stimulation, and then amplifier was changed back to current clamp to assess membrane properties in the presence of hymeglusin. After returning to voltage clamp and restarting rhythmic stimulation, 2DG-0G was applied in the presence of hymeglusin for the next 30 minutes. In a subset of hibernator cells in the presence of hymeglusin, n=7 continued to be rhythmically stimulated with 2DG-0G for the next 30 minutes (60 minutes in total) or n=8 had 5 mM BHB added during the next 30 minutes while being stimulated in the presence of 2DG-0G (60 minutes in total, first 30 minutes 2DG-0G+hymeglusin and the last 30 minutes 2DG-0G+hymeglusin+BHB). In a different set of experiments, control neurons (n=15) were exposed to the rhythmic stimulation protocol in the presence of 2DG-0G with 5 mM BHB.

Finally, to assess the involvement of astrocytes, we incubated slices for 2 hr in 1 mM L-α-aminoadpic acid (AAA), as this treatment has been demonstrated to induced death in astrocytes while having minimal effects on neurons(13-15). While we are not aware of studies tracing the developmental lineage of astrocytes in frogs, the central nervous system of frogs contains glial cells consistent with astrocytes, with glycogen granules(16), Glial Fibrillary Acidic Protein (GFAP) expression, and an ohmic $K^+$ selective membrane(17). Following incubation of slices, the same protocol was performed as before with 2DG-0G during rhythm stimulation of EPSCs. In n=3 cells, AAA was applied acutely for 30 minutes and did not influence synaptic transmission. In all experiments, negative pressure was applied to the microelectrode to maintain series resistance through the experimental period. If series resistance changed by 20% and could not be reduced via negative pressure, the experiment was abandoned. One slice was used per exposure to 2DG-0G. If the recording was lost in the middle of 2DG-0G, the slice was discarded.

All patch clamp electrophysiology data were analyzed using the Easy Electrophysiology data analysis program (Easy Electrophysiology Ltd., London, UK). Synaptic transmission was assessed as the area under the curve (AUC) during 1 minute of stimulation at the end of each sampling period. Firing rate was taken as the firing rate during 500 ms current injections and is presented as either maximally firing rate or the entire frequency vs. current relationship. Input resistance is taken as the change in voltage change from resting membrane potential during a 100 pA step.

**Drugs**
DNQX was from Hello Bio (Princeton, NJ, USA). Hymeglusin was from Cayman Chemical (Ann Arbor, MI, USA). L-α-aminoadpic acid (AAA) was from Thermo Scientific Chemicals (Waltham, MA, USA). R)-(-)-3-Hydroxybutyric acid sodium salt (BHB) was Santa Cruz Biotechnology Inc. (Dallas, TX, USA).

**Tissue BHB determination**

To determine the influence of inhibiting glucose metabolism on tissue beta hydroxybutyrate (BHB) we used an experimental approach that allows each brain to serve as its own control. Following dissection of the brainstem-spinal cord in controls and hibernators as described above, the preparation was transected down the midline, separating the two hemispheres. One half was left in normal aCSF (control half), and the other was exposed to 2DG-0G to inhibit glucose metabolism (2DG-0G half). After one hour in 2DG-0G, 40 mg of brainstem from each half was quickly dissected, dried of excess liquid, weighed, minced into small pieces, and then added to separate 1.5 mL tubes containing 229 µL of Tris buffer. The contents were then homogenized with a mechanical tissue homogenizer. To measure tissue BHB, we used a beta hydroxybutyrate colorimetric assay kit from Cayman Chemical following the manufactures instructions (kit #700190, Ann Arbor, MI, USA). Briefly, the homogenate was centrifuged at 1,000 x *g* for 10 minutes at 4°C. The supernatant was removed and then added to a microcentrifuge tube and centrifuged at 10,000 x *g* for 10 minutes at 4°C. The supernatant was removed and the pellet was resuspended in 150 µl of assay buffer. Resuspended pellet was then stored in a -80 freezer until all samples were collected. BHB concentrations were assayed in all samples run in duplicate on the same plate using a Varioskan LUX plate reader (Thermo, Waltham, MA, USA) and quantified by constructing a standard curve the following manufacturer's instructions. We then normalized BHB concentrations to total protein from each sample in duplicate using a Bradford Assay. The BHB concentration for each 2DG-0G half was divided by the concentration from the control half to determine the change in BHB in response to 2DG-0G within the same brain.

**RNA isolation and real-time quantitative PCR**

All methods for sample collection, RNA isolation, cDNA synthesis of 1 µg of cDNA, and real-time quantitative PCR (qPCR) using 1/5 diluted cDNA template were performed as described previously by our lab(18). This study, we identified candidate genes involved in steps of beta oxidation, ketogenesis, ketolysis, and BHB transport (Table 1). We first sought to identify annotated genes in the *Aquarana catesbeiana* genome. Only 1 candidate gene was annotated. We then used sequences from *Rana temporaria* or *Xenopus laevis*, frog species with well-annotated genomes, to query the coding sequence of non-annotated "hypothetical proteins" in the *Aquarana catesbeiana* genome(19). Once a hypothetical protein was identified as likely to be homologous to that of *Rana temporaria* or *Xenopus laevis*, the animo acid sequence was BLASTed against the entire non-redundant protein data base (BLAST, NCBI, USA). In the case of 6 candidate genes, queries returned hits of high sequence identity with the excepted results. For example, a protein BLAST of *Aquarana catesbeiana* HMGCS1 returned a list of hits containing HMGCS1 from a variety of other species. We then used the coding DNA sequence (CDS) to design qPCR primers for these candidates. If we were unable to identify annotated or hypothetical proteins with relevant hits in the reBLAST of the sequence from *Aquarana catesbeiana*, we used CDS sequences from *Rana temporaria* to design qPCR primers, as this species is closely-related containing a genome with a chromosome-level assmebly(20). All qPCR primers were designed using PrimerBlast and validated in-house. For this, all primers were tested using a 4-fold dilution series and produced a single peak during melt curve analysis. If these criteria were not met, forward and reverse primers were redesigned. Changes in gene expression were determined using the $2^{-\Delta\Delta ct}$ method(21). 18s ribosomal RNA (rRNA) from stock cDNA (original 1 µg diluted 5-fold) was further diluted by 50-fold and used as the control gene in analysis. $C_t$ values of 18S rRNA for controls (20.4±0.4) and hibernators (20.3±0.5) were not different (p=0.93, two-tailed unpaired t test).

#### Table 1. Primer information for candidate genes

| Targets | Forward Primer Sequence | Reverse Primer Sequence | PCR Efficiency (%) | NCBI accession number *Aquarana catesbeiana* | NCBI accession number *Rana temporaria* |
|---|---|---|---|---|---|
| **CPT-1L** | GGGCCATCTCTGGGAGTACA | CAGTCTGCTTGGAGGAGGGA | 91.6 | PIO37846.1 | ▪ |
| **CPT-1M** | AGCTGGGGACCGAGTTTCTT | GCATCAGGGTCCAGGCAAAC | 96.4 | PIO31695.1 | |
| **CPT2** | GGTGCAGCTAGGAGGCTTTG | GCAACCAATCCAGTCGTCGT | 94.1 | ▪ | XM_040360458.1 |
| **LCAD** | GACAGGTGAGCCGGGAATTG | CACCAATGCCACCATGCTCT | 94.2 | ▪ | XM_040357444.1 |
| **VLCAD** | TCCGGCTGTTTGTTGCTCTC | GCTCAGTGGATTGGTGACGG | 90.5 | ▪ | XM_040334301.1 |
| **MCAD** | ACGCGATGGAGAGGAAGACA | CCCAGGCTGCTCTCTGGTAT | 90.4 | ▪ | XM_040360675.1 |
| **ACAA1** | GGACGTCAACCTGAAGCCTG | GGCAGCTCTAGCAACCACAG | 94.7 | ▪ | XM_040353382.1 |
| **ACAA2** | CACTGCTGAGAACCTGGCTG | GTCCTGAGCGGCTTTCCATC | 90.9 | ▪ | XM_040336875.1 |
| **HMGCS1** | CCTCAGGGCTCAGTGGATGA | GCCTCCAGGGGTCCATCATT | 98.6 | PIO24663.1 | ▪ |
| **HMGCL** | ACCTGTGCGAGGGTATGTGT | TGTGTGCCACCTCTGCTACT | 90.0 | ▪ | XM_040343136.1 |
| **BDH1** | CCAAGGCCTGTCTCCCTCTT | GGGGGAACGAGCTGGATTTG | 98.9 | PIO23142.1 | ▪ |

| | | | | | |
|---|---|---|---|---|---|
| **BDH2** | GACACGAGTGCTGGATGTGA | GGATCCGTGGTGGACAAAGC | 85.0 | ACO51978.1 | |
| **ACAT2** | TTGTGGTTGCCGGTGGTATG | TGGGACATCCCCAGCTTTCA | 94.1 | ▪ | XM_040340584.1 |
| **OXCT1** | AGCGTCATTCTTCTCCAGCG | TTGACACCTGCATTGCTCCC | 97.6 | ▪ | XM_040338845.1 |
| **AACS** | GGGAGTGGAAGCATGGTGTG | TGTGTGGGCTGAGATGGGAG | 97.5 | PIO41121.1 | ▪ |
| **MCT1** | TGGCTTTGCTTTTGGCTGGC | GGCCCAAGAAGAACAGGGCA | 90.0 | ▪ | XM_040337478.1 |
| **MCT2** | TGCCAGGCCGTTAACTGGAA | GACAAGGCCCGCGTAGGAAG | 93.4 | PIO23679.1 | ▪ |
| **MCT4** | CCCCGTCTTCCTCTGTTGCC | ATAAGAAAGCCACCGCGCCA | 88.0 | ▪ | 040330784.1 |
| **18S rRNA** | CAGGCCGGTCGCCTGAATAC | GGCCCCAGTTCCGAAAACCA | 101.068 | KV934453.1 | ▪ |

**Statistics and data presentation**

Data are presented as mean ± standard deviation or box and whisker plots, with single data points from individual experiments, where it is most appropriate for visualization. For physiology assessments and measurements of tissue BHB, when data sets were approximately normally distributed, parametric statistical tests were used. When two groups were compared in before-after/time course experiments (i.e., with dependent samples), a two-tailed paired t-test or repeated measures two-way ANOVA were used. When groups contained independent samples, a two-tailed unpaired t-test or one-way ANOVA was used. One-way and two-way ANOVAs were followed with Holm-Sidak multiple comparisons tests. In cases where data sets were not normally distributed, non-parametric statistical tests were used. In these cases, when analyzing two independent groups, a Mann-Whitney tests was used, and when three with three or more groups, a Kruskal-Wallis test was used. For mRNA expression, all comparisons for each of the 18 candidate genes between controls and hibernators were compared two-stage linear step up procedure of Benjamini, Krieger,and Yekutieli to control for the false discovery rate. Significance was accepted when $p< 0.05$. All analyses were performed using GraphPad Prism (v9.4.1, San Diego, CA, USA).

**Supplemental Figures**

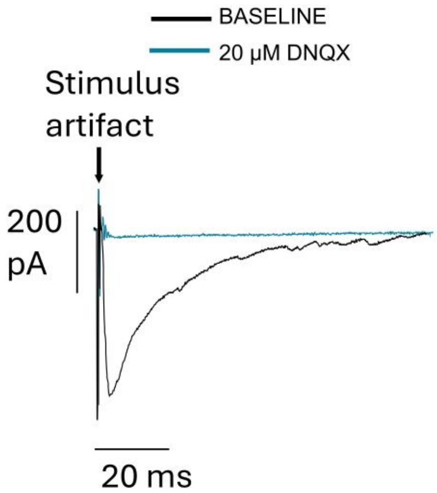


**Supplemental Figure 1 Stimulation of solitary tract axons evokes excitatory postsynaptic currents carried by AMPA-glutamate receptors on hypoglossal motoneurons.** Axon stimulation while voltage clamping motoneurons at -80 mV leads to inward currents that are blocked by 20 µM DNQX. This example is representative of 5 different cells from 3 animals where DNQX abolished evoked EPSCs.

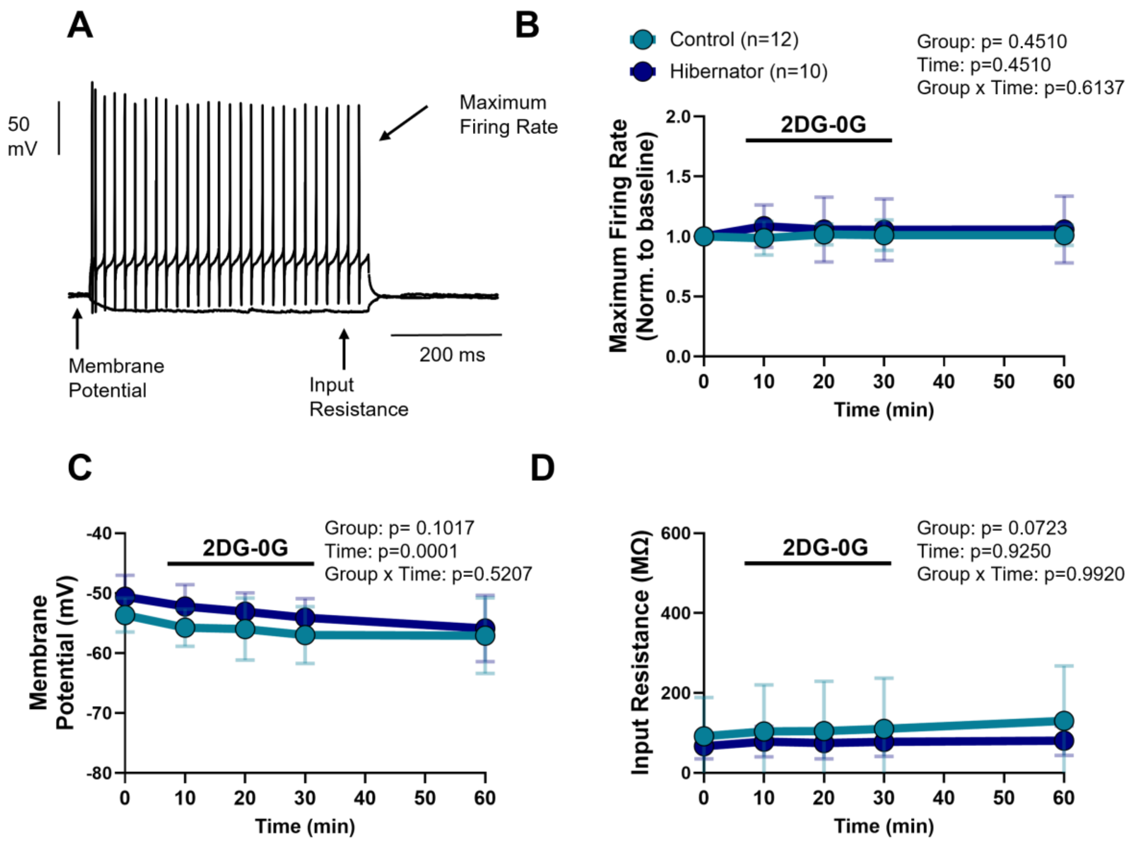


**Supplemental Figure 2 Intrinsic membrane properties respond the same in controls and hibernators in 2DG-0G.**

A) Example current clamp recording of membrane potential during current injections to address intrinsic membrane properties.

B) Mean±SD of maximum firing in these experiments evoked by 1 nA current injection to the cell body normalized to baseline. 2DG-0G exposure did not influence maximum firing rate. Control, n=12 neurons from N= 5 animals. Hibernators, n=10 cells from 5 animals. Results from repeated measures two-way ANOVA shown in graph.

C) Mean±SD of resting membrane potential. 2DG-0G exposure slightly hyperpolarized membrane potential over the course of the 60 min experiment (significant time effect in repeat measures two-way ANOVA). However, no group or time x group interaction, indicating this occurred the same in controls and hibernators. Although this change was statistically significant, the effect was small. Controls membrane potential changed -3.4±5.0 mV and hibernators -5±4.50 mV from the start to the end of the experiment. Control, n=12 neurons from N= 5 animals. Hibernators, n=10 cells from 5 animals. Results from repeated measures two-way ANOVA shown in graph.

D) Mean±SD of input resistance (change in membrane potential divided by the current step). 2DG-0G exposure did not influence input resistance. Control, n=12 neurons from N= 5 animals. Hibernators, n=10 cells from 5 animals. Results from repeated measures two-way ANOVA shown in graph.

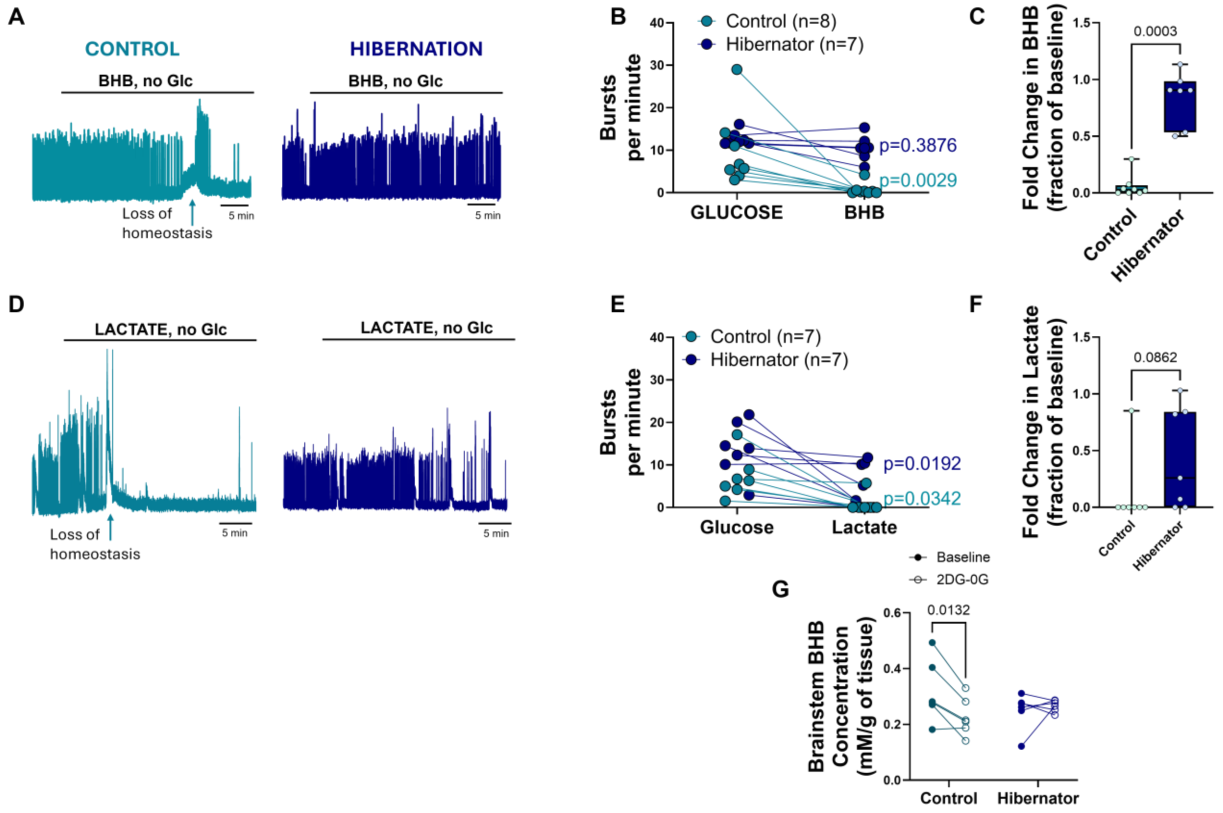


**Supplemental Figure 3 BHB substitution supports network activity better than lactate after hibernation.** (A) Example recording of network activity recorded through cranial motor nerves in control and hibernator during equimolar substitution of BHB for glucose. In controls, fuel substitution leads to a loss of homeostasis seen as large increases in background activity, and then a loss of rhythmic activity. Hibernators persist unabated when BHB is substituted for glucose. (B-C) Plot shows absolute burst rates before and after the substitution of BHB for glucose. Repeated measures two-way ANOVA results: group, p=0.0042, fuel, p=0.0048, and group x fuel interaction, p=0.0567. P values shown on the graph are from Holm-Sidak multiple comparisons tests indicating that controls significantly decrease activity when only using BHB, while hibernators continue normal activity. The graph on the right is the same data plotted with burst rate in BHB relative to activity with glucose (p value from a Mann-Whitney U test). (D) Example recording of network activity recorded through cranial motor nerves in control and hibernator during the substitution of lactate (15 mM) for glucose. In controls, fuel substitution leads to a loss of homeostasis seen as large increases in background activity, and then a loss of rhythmic activity. Hibernators slow over time with lactate substitution. (E-F) Plot shows absolute burst rates before and after the substitution of lactate for glucose. Repeated measures two-way ANOVA results: group, p=0.0096, fuel, p=0.0023, and group x fuel interaction, p=0.6373. P values shown on the graph are from Holm-Sidak multiple comparisons tests indicating that controls and hibernators significantly decrease activity when only using lactate. The graph on the right is the same data plotted with burst rate in lactate relative to activity with glucose (p value from a Mann-Whitney U test). (G) Absolute tissue BHB concentrations from split brain preparations with one half held in regular aCSF and the other half in 2DG-0G for 60 minutes. Controls decrease while hibernators maintain BHB concentrations (repeated measures two-way ANOVA- significant interaction between group x treatment, p=0.0163; Holm-Sidak multiple comparisons tests, only significant difference for control baseline vs. 2DG-0G).

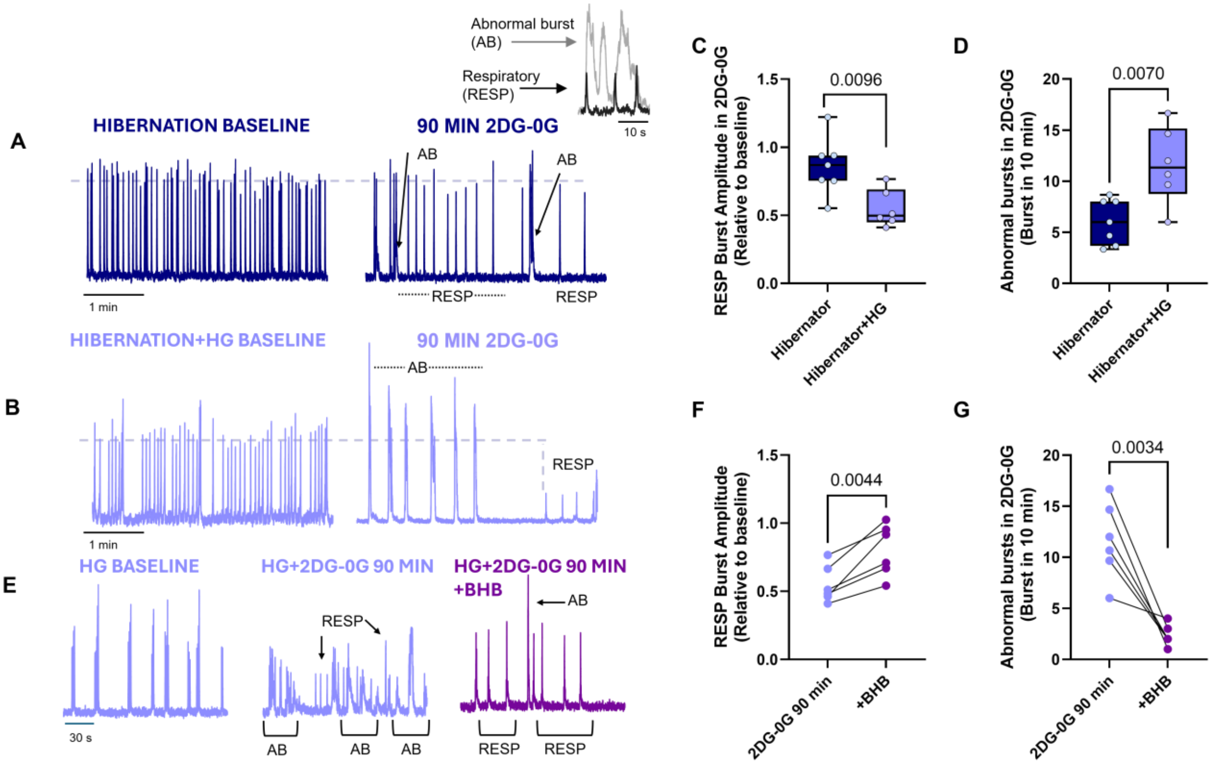


**Supplemental Figure 4 Brain-derived ketone bodies support network activity without glucose metabolism**

A-B) Example recordings of rhythmic motor nerve recordings from hibernator preparations without (A) and with 2.5 µM hymeglusin (B) at baseline and after 90 minutes of 2DG-0G. Rhythmic motor activity in intact circuits was well maintained for at least 90 minutes in 2DG-0G, as indicated by the stable amplitude of the rhythmic bursts. In contrast, hibernators exposed to hymeglusin still exhibited motor output but normal respiratory-related bursts were smaller in amplitude at 90 minutes, corroborating weakened, but still functional, synaptic transmission that reduces motor recruitment. Chaotic seizure-like activity was more prominent during 2DG-0G in hibernators with hymeglusin, indicative of a generalized loss of homeostasis in the brainstem without ketone body synthesis. Inset shows the distinction between respiratory and abnormal, non-respiratory burst types.

C) Box and whisker plot showing burst amplitude at the end of 90 minutes of 2DG-0G without (hibernator) and with hymeglusin (hibernator+HG). P value is from unpaired two-tailed t test. N=7 hibernators and N=6 hibernator+hymeglusin

D) Box and whisker plot showing the frequency of non-respiratory, chaotic activity the end of 90 minutes of 2DG-0G without (hibernator) and with hymeglusin (hibernator+HG). P value is from unpaired two-tailed t test.

E) To verify that these decrements in function were caused by hymeglusin's ability to disrupt ketone body biosynthesis, 5 mM BHB was bath-applied following 90 minutes of 2DG-0G+hymeglusin. Example recordings of rhythmic motor nerve recordings from hibernator preparations baseline with hymeglusin (left), after 90 minutes of 2DG-0G (middle), and then after restoring BHB (right). Supplementing BHB after inhibiting BHB synthesis increased normal burst amplitude and reduced non-respiratory, seizure-like activity.

F) Individual responses of burst amplitude (relative to baseline) at the end of 90 minutes of 2DG-0G and 2.5 μM hymeglusin and then after providing 5 mM BHB. Burst amplitude increased in each experiment. P value is from two-tailed paired t test. N=6

G) Individual responses of non-respiratory, chaotic activity at the end of 90 minutes of 2DG-0G and 2.5 μM hymeglusin and then after providing 5 mM BHB. Burst amplitude increased in each experiment. P value is from two-tailed paired t test. N=6

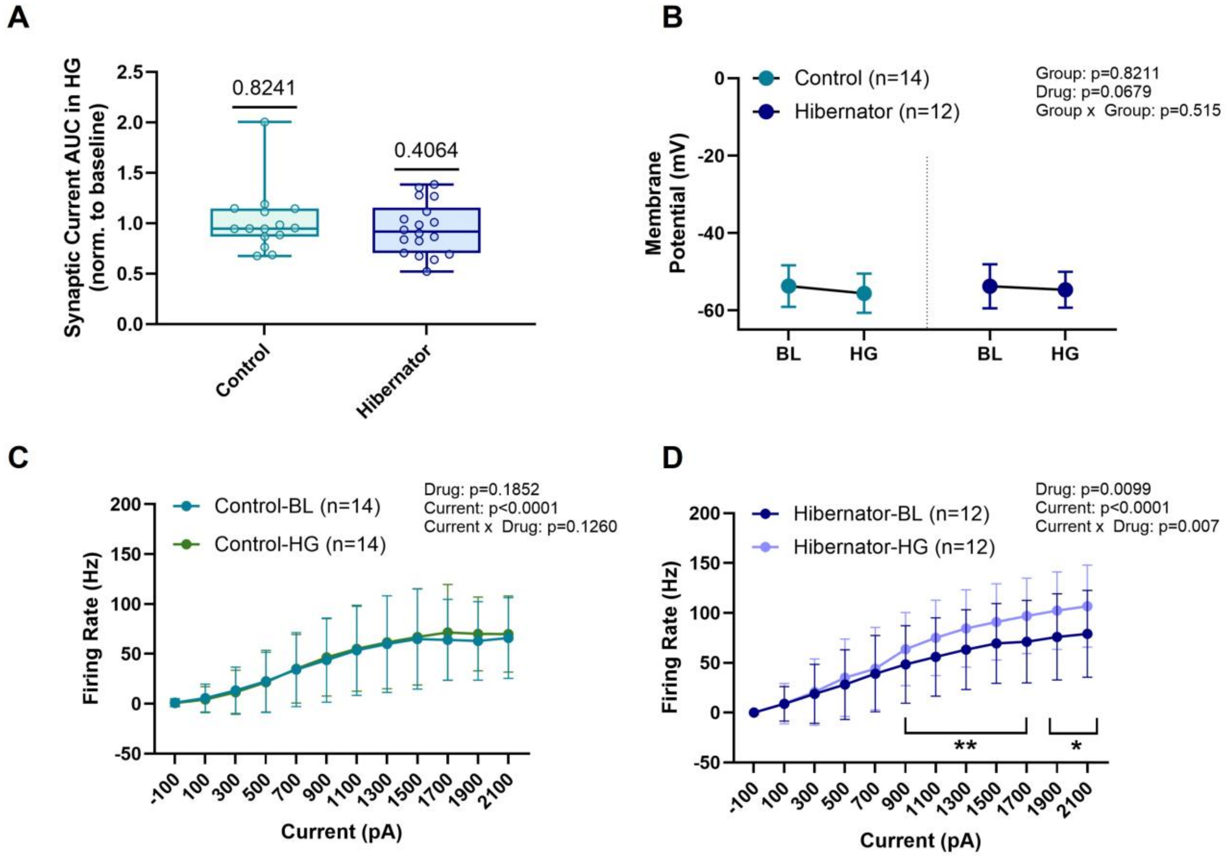


**Supplemental Figure 5 Influence of hymeglusin on baseline neuronal properties when glucose metabolism is intact.**

A) Box and whisker plots showing the change in rhythmically evoked synaptic transmission (area under the curve) in response to 2.5 μM hymeglusin. P values shown are from one-sample t tests, indicating that hymeglusin does not change synaptic activity from baseline levels on average. Comparing the two groups with a two-tailed unpaired t test yields a p value of 0.4874, indicating there are also no differences between groups. Control, n=15 cells from N=5 animals. Hibernator, n=18 cells from N=8 animals.

B) Mean±SD of membrane potential before and after application of hymeglusin. Hymeglusin does not influence resting potential in controls and hibernators. Repeated measure two-way ANOVA.

C-D) Mean±SD of hypoglossal motoneuron firing rate vs. input current stimulus before (baseline, BL) and after hymeglusin (HG). C) control, D) hibernator. Hymeglusin does not affect firing rates in controls but increases firing rate in hibernators at higher current intensities. Repeated measures two-way ANOVA outputs are presented in the plots. ** indicates p<0.01 and *p<0.05 from Holm-Sidak Multiple comparisons test.

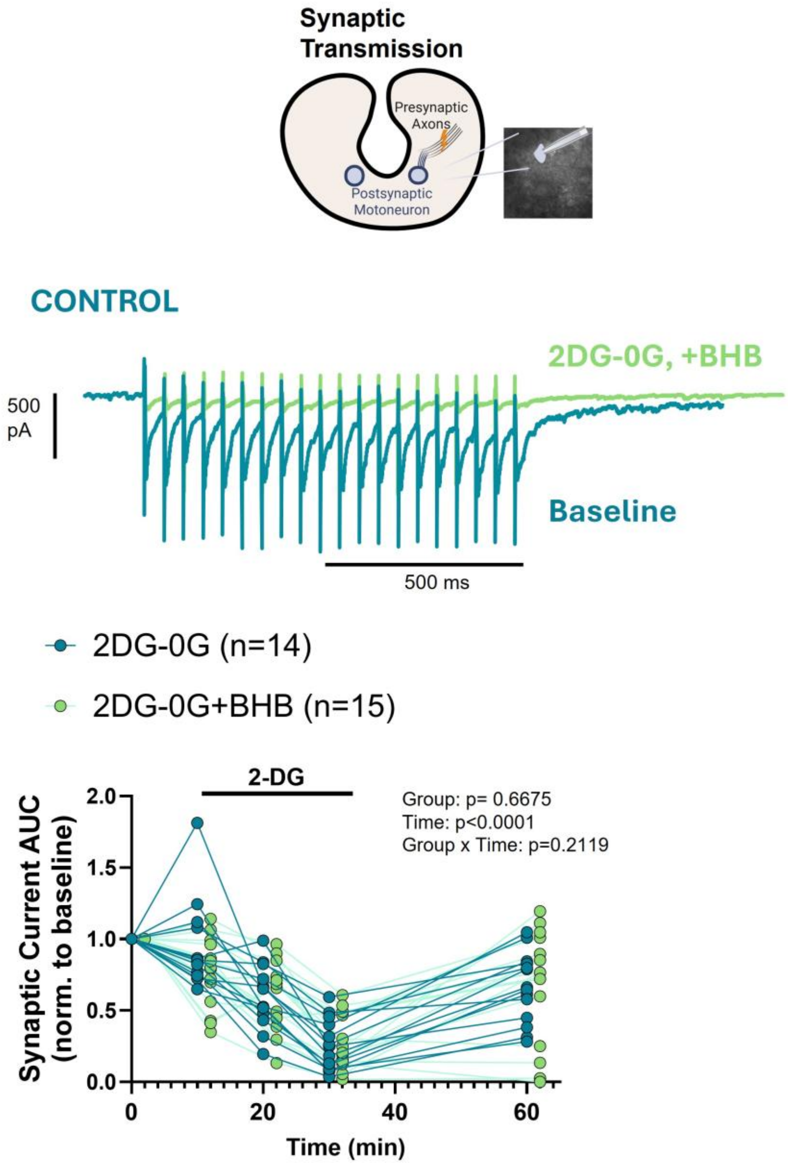


**Supplemental Figure 6 BHB as a lone fuel cannot support synaptic and network activity in control animals.** Example recordings of rhythmically-evoked synaptic transmission in brainstem slices in controls with 2DG-0G and with 2DG-0G+5 mM BHB (top). On the bottom shows data for synaptic transmission from individual experiments from controls with 2DG-0G (n=14 from N=6 animals) and 2DG-0G+BHB (n=15 from 4 animals). The time course of the decrease in was the same and was not influenced by adding 5 mM BHB in controls. Repeated Measures two-way ANOVA shown in plot.

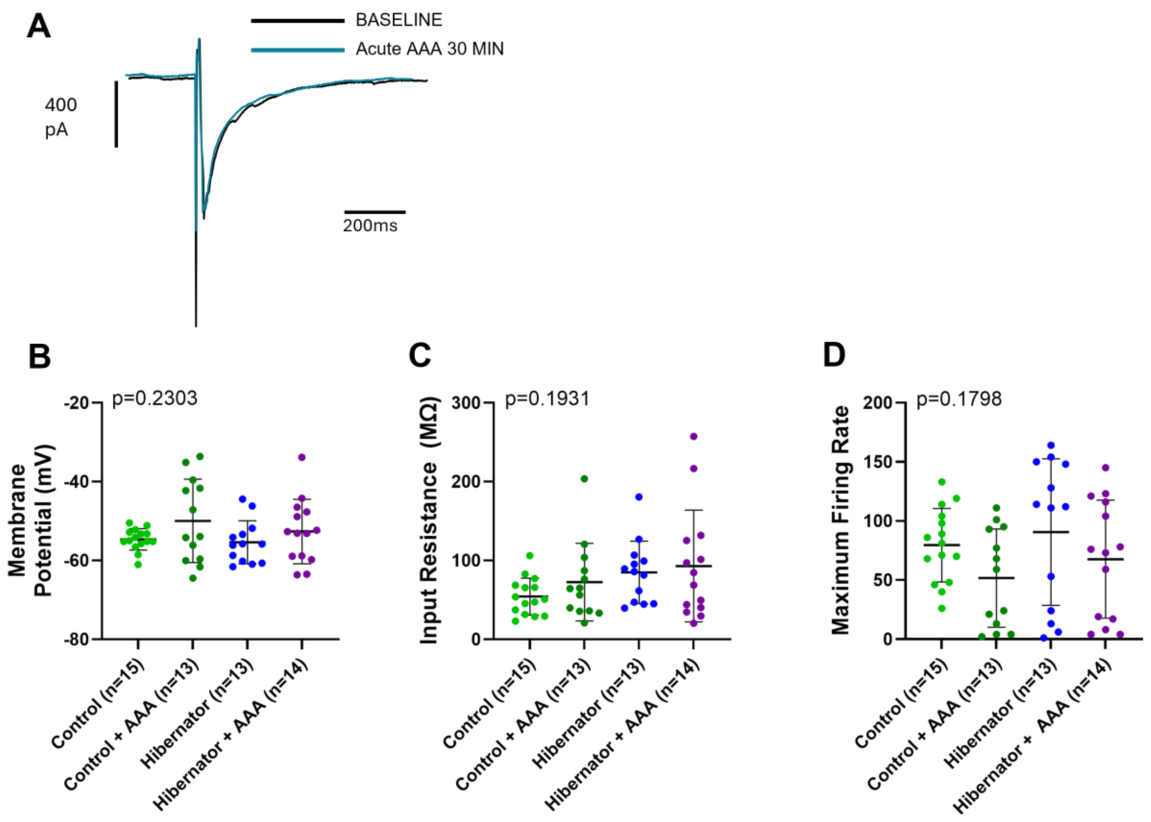


**Supplemental Figure 7 L-α-aminoadpic acid (AAA) does not influence baseline neuronal properties.**

A) Example recording of evoked synaptic transmission before and after a 30 minute exposure to 1 mM AAA in a control slice. Experiment is representative of 3 different cells with similar results.

B-D) 2 hr incubation of slices with 1 mM AAA does not influence membrane potential (B; p value from one-way ANOVA), input resistance (C; p value from Kruskal-Wallis test), and maximum firing rates (D; p value from Kruskal-Wallis test) in controls and hibernators.

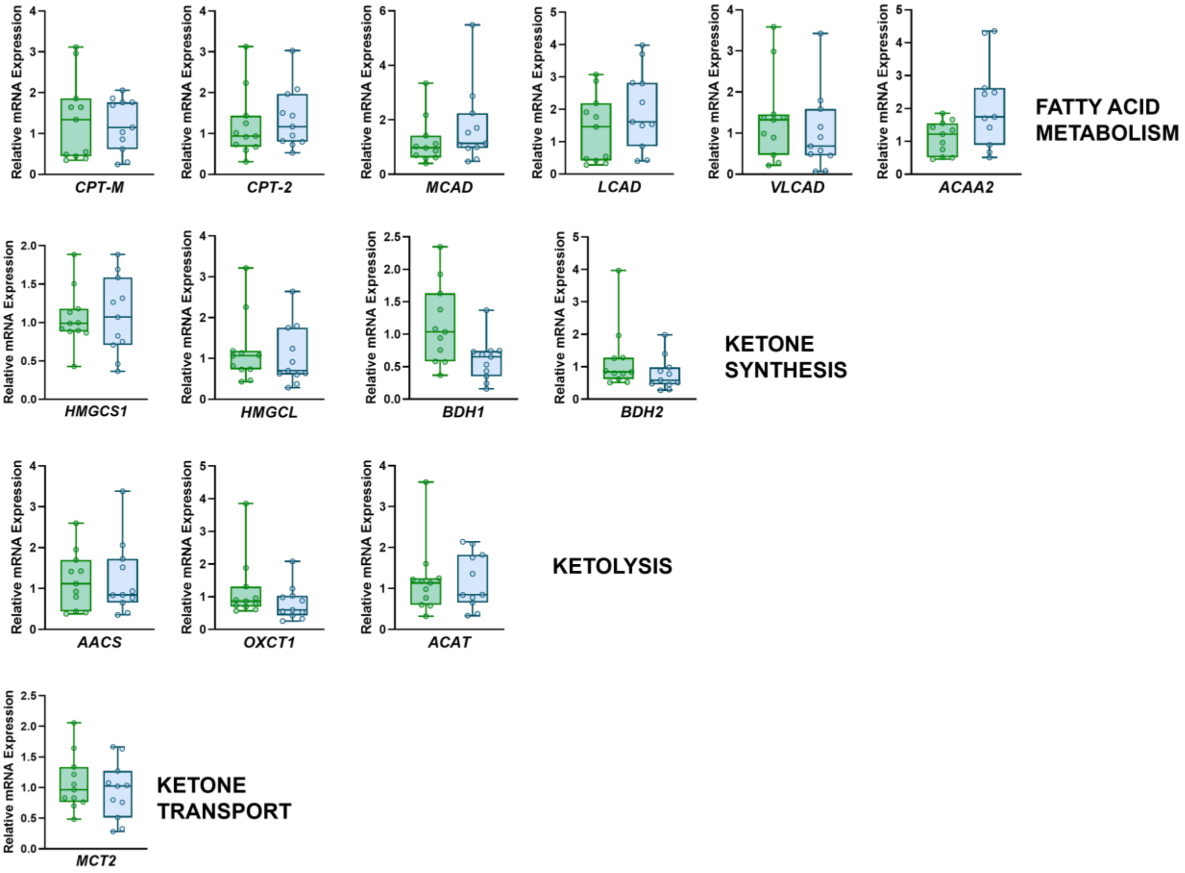


**Supplemental Figure 8 Candidate genes involved in fatty acid metabolism, ketogenesis, ketolysis, and beta hydroxybutyrate transport that did not change expression.** CPT-M (carnitine palmitoyl transferase-muscle), CPT2 (carnitine palmitoyl transferase 2), MCAD (medium chain acyl-CoA dehydrogenase), LCAD (long chain acyl-CoA dehydrogenase), VLCAD (very long chain acyl-CoA dehydrogenase), ACAA2 (acetyl-CoA acyltransferase 2), HMGCS1 (3-Hydroxy-3-methylglutaryl-CoA synthase 1), HMGCL (3-hydroxy-3-methylglutaryl-CoA lyase), BDH1 (3-hydroxybutyrate dehydrogenase 1), BDH2 (3-hydroxybutyrate dehydrogenase 2), AACS (acetoacetyl-CoA synthetase), OXCT1 (3-oxoacid CoA transferase 1), ACAT (Acetyl-CoA acetyltransferase), MCT2 (monocarboxylate transporter 2). q values from the two-stage linear step up procedure of Benjamini, Krieger,and Yekutieli to control for the false discovery rate all >0.05.  n=11 controls and n=11 hibernators per analysis.

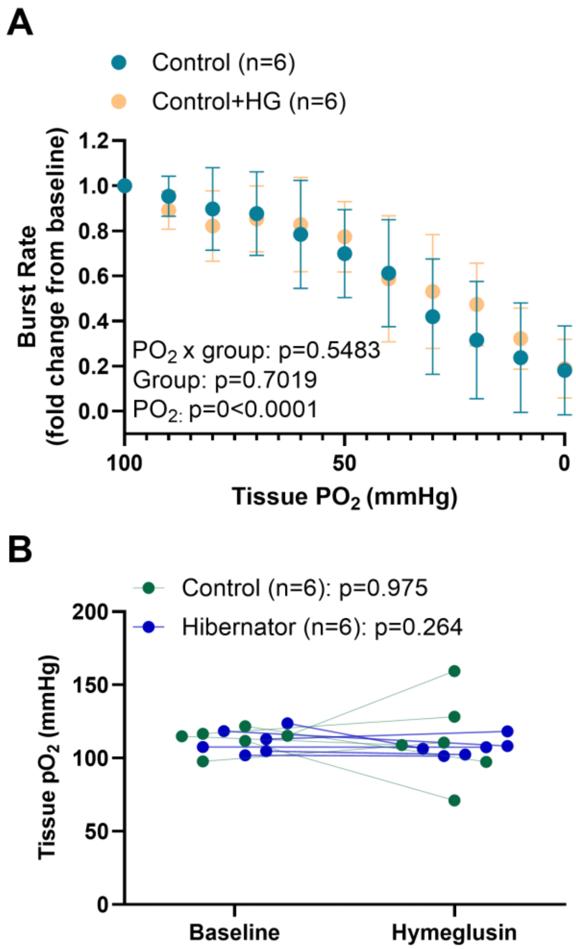


**Supplemental Figure 9 Hymeglusin does not influence hypoxia-sensitivity in controls and does not alter apparent baseline $O_2$ consumption in controls and hibernators.**

A) Mean±SD of network burst rate vs. tissue $pO_2$ in control animals with and without 2.5 μM hymeglusin (HG). Outputs from repeated measures two-way ANOVA are presented in the plot.

B) Tissue $pO_2$ before and after application of 2.5 μM hymeglusin. In controls and hibernators, hymeglusin did not influence tissue $pO_2$. p values are from paired t tests.
